# No evidence for prolactin’s involvement in the post-ejaculatory refractory period

**DOI:** 10.1101/2020.08.19.257196

**Authors:** Susana Valente, Tiago Marques, Susana Q. Lima

## Abstract

In many species, ejaculation is followed by a state of decreased sexual motivation, the post-ejaculatory refractory period. Several lines of evidence have suggested prolactin, a pituitary hormone released around the time of ejaculation in humans and other animals, to be a decisive player in the establishment of the refractory period. However, data supporting this hypothesis is controversial. We took advantage of two different strains of house mouse, a wild derived and a classical laboratory strain, that differ substantially in their sexual behavior, to investigate prolactin’s involvement in sexual motivation and the refractory period. First, we show that there is prolactin release during sexual behavior in male mice. Second, using a pharmacological approach, we show that acute manipulations of prolactin levels, either mimicking the natural release during sexual behavior or inhibiting its occurrence, do not affect sexual motivation or shorten the refractory period, respectively. Therefore, we show compelling evidence refuting the idea that prolactin released during copulation is involved in the establishment of the refractory period, a long-standing hypothesis in the field of behavioral endocrinology.

## Introduction

Sexual behavior follows the classical sequence of motivated behaviors, terminating with an inhibitory phase after ejaculation: the post-ejaculatory refractory period (PERP)^1^. The PERP is highly conserved across species and includes a general decrease in sexual motivation and also inhibition of erectile function in humans and other primates^2^. This period of time is variable across and within individuals and is affected by many factors, such as age^3,4^ or the presentation of a new sexual partner^5,6^. The PERP is thought to allow replacement of sperm and seminal fluid, functioning as a negative feedback system where by inhibiting too-frequent ejaculations an adequate sperm count needed for fertilization is maintained^7,8^.

Several lines of evidence have suggested the hormone prolactin (PRL) to be a key player in the establishment of the PERP^9,10^. PRL is a pleiotropic hormone, first characterized in the context of milk production in females, but for which we currently know several hundred physiological effects in both sexes^11,12^. The association of PRL to the establishment of the PERP in males is based on several observations. First, it was shown that PRL is released around the time of ejaculation in humans and rats^13–21^. Anecdotally, no PRL release has been observed in a subject with multiple orgasms^22^. Second, chronically abnormal high levels of circulating PRL are associated with decreased sexual drive, anorgasmia and ejaculatory dysfunctions^23,24^. Finally, removal of PRL-producing pituitary tumors or treatment with drugs that inhibit PRL release reverse sexual dysfunctions^25,26^. Taking these observations into consideration, it has been hypothesized that the PRL surge around the time of ejaculation plays a role in the immediate subsequent decrease of sexual arousal, the hallmark of the PERP. In fact, this idea is widespread in behavioral endocrinology textbooks^27^ and the popular press^1^*.

PRL is primarily produced and released into the bloodstream from the anterior pituitary^11,28^, reaching the central nervous system either via circumventricular regions lacking a blood-brain barrier^29^ or via receptor-mediated mechanisms^30^, binding its receptor which has widespread distribution, including in the social brain network^31^. Hence, circulating PRL can impact the activity of neuronal circuits involved in the processing of socio-sexual relevant cues and in principle alter the detection of opposite-sex conspecific cues and thus sexual arousal^32,33^. Circulating PRL reaches the central nervous system on a timescale that supports the rapid behavioral alterations that are observed immediately after ejaculation (in less than 2 minutes)^34^. Through mechanisms that are not yet well established, PRL elicits fast neuronal responses^35^ besides its classical genomic effects^36^. In summary, circulating PRL can reach the brain and affect brain regions involved in socio-sexual behavior on a time scale compatible with the establishment of the PERP.

However, despite data supporting the involvement of the ejaculatory PRL-surge in the establishment of the PERP, this hypothesis has received numerous critics^2,3,37–39^. While in humans it is well established that chronically high levels of PRL reduces sexual motivation^24^, some authors suggest that those results were erroneously extended to the acute release of PRL^2,3,37–39^. Furthermore, there is controversy in relation to PRL dynamics during sexual behavior, since in most studies PRL levels were quantified during fixed intervals of time, and not upon the occurrence of particular events, such as ejaculation. In fact, some reports in rats suggest that PRL levels are elevated through the entire sexual interaction^40,41^. Finally, formal testing of the impact of acute PRL manipulations on sexual motivation and performance is still missing (but see^42^ for an acute manipulation in humans).

In the present study, we tested the role of PRL in sexual motivation and in the establishment of the PERP in the mouse. The sequence of sexual behavior in the mouse is very similar to the one observed in humans^43^, making it an ideal system to test this hypothesis. Also, we took advantage of two strains of inbred mice that are representative of two different mouse subspecies (C57BL/6J: laboratory mouse, predominantly *Mus musculus domesticus* and PWK/PhJ: inbred wild-derived, *Mus musculus musculus*^44^) and exhibit different sexual performance. Through routine work in our laboratory, we observed that while most BL6 males take several days to recover sexual interest after ejaculation, a large proportion of PWK males will re-initiate copulation with the same female within a relatively short period of time. This difference in PERP duration can be taken to our advantage, widening the dynamic range of this behavioral parameter and increasing the probability of detecting an effect of the manipulation.

By monitoring PRL levels in sexually behaving male mice and pharmacological manipulations, we specifically asked the following questions: (i) what is the PRL release dynamics during sexual behavior? (ii) is an acute PRL release sufficient to decrease sexual motivation, the hallmark of the PERP? And consequently (iii) does blocking the acute release of PRL during copulation shorten the duration of the PERP?

## Results

### Prolactin is released during sexual behavior in male mice

We first asked if PRL is released during copulation in our two strains of male mice. To monitor PRL dynamics during sexual behavior we took advantage of a recently developed ultrasensitive ELISA assay that can detect circulating levels of PRL in very small volumes of whole blood (5-10 microliters), allowing the assessment of longitudinal PRL levels in freely behaving mice^45^. Sexually trained laboratory mice (C57BL/6J, from here on BL6) or inbred wild-derived mice (PWK/PhJ, from here on PWK) were paired with a receptive female and allowed to mate (see Methods for details). During the sexual interaction males were momentarily removed from the cage to collect tail blood after which they returned to the behavioral cage, resuming the sexual interaction with the female. We collected blood samples upon the execution of pre-determined, easily identifiable, behavioral events that correspond to different internal states of the male: before sexual arousal (*baseline*, before the female was introduced in the cage), at the transition from appetitive to consummatory behavior (*mount attempt*, immediately after the male attempted to mount the female for the first time), during consummatory behavior (*mount*, after a pre-determined number of mounts with intromissions, BL6=5 and PWK=3) and immediately after ejaculation (*ejaculation*, after the male exhibited the stereotypical shivering and falling to the side) (Fig.1a, please see Methods for details).

**Fig. 1.**
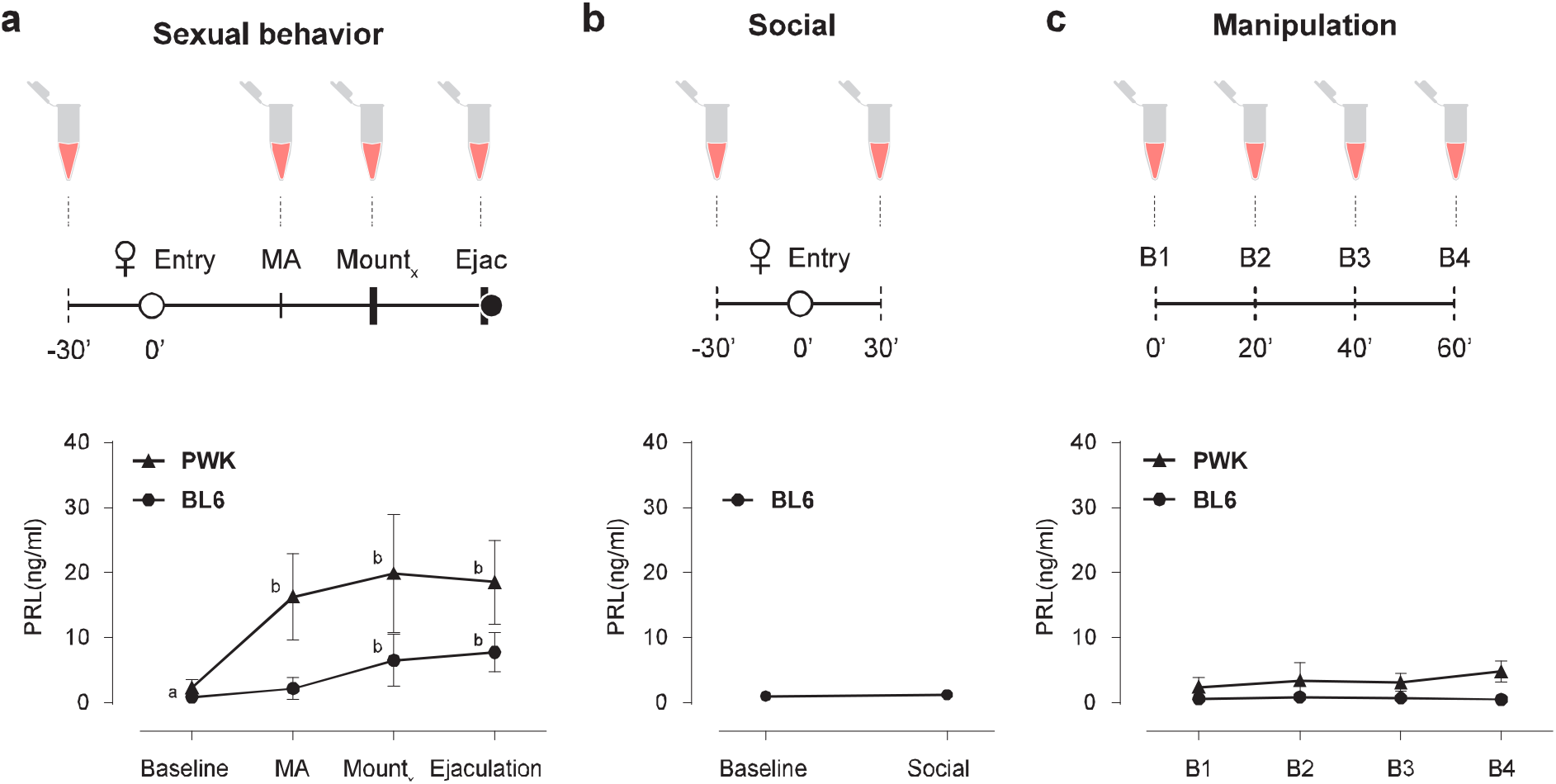
Prolactin is released during sexual behavior in male mice. **a** Timeline for blood collection and [PRL]_blood_ during sexual behavior (MA-mount attempt; BL6_Xmounts_ = 5; PWK_Xmounts_ = 3). RM One-way Anova for BL6 (n = 8) F_3,7_=21.26, *P* < 0.0001 and PWK (n = 9) F_3,8_=17.18, *P* < 0.0001, followed by Tukey’s multiple comparison test ab *P* ≤ 0,01. **b** Timeline for blood collection and [PRL]_blood_ during social behavior in BL6 males (n = 15) *P* = 0.282, two-tailed Paired *t* test. **c** Timeline for blood collection and [PRL]_blood_ during repeated sampling in resting condition. RM One-way Anova for BL6 (n = 8) F_3,7_ = 2.08; *P* = 0.18 and PWK (n = 8) PWK F_3,7_ = 2.94; *P* = 0.11. Data represented as mean ± SD.

Baseline levels of circulating PRL in male mice were low for both strains (BL6 0.86 ± 0.46; PWK: 2.31 ± 1.37 ng/ml; please see^45^ for BL6), but are significantly increased during sexual interaction (Bl6: F_3,7_ = 21.26; *P* < 0.0001; PWK F_3,8_ = 17.18; *P* < 0.0001, RM One-way ANOVA (Fig. 1a). While in the case of BL6 males PRL levels only increased during the consummatory phase, PRL levels in PWK males are significantly increased already at the transition from appetitive to consummately behavior (baseline *vs* MA 16.30 ± 6.67 ng/ml, *P* = 0.001, Tukey’s multiple comparisons test) (Fig. 1a). In both strains, PRL levels after ejaculation are similar to the levels reached during consummatory behavior (BL6 *P* = 0.71 vs PWK *P* = 0.95, Tukey’s multiple comparisons test), in marked contrast to humans, where PRL seems to be released only around the time of ejaculation^15^.

Contrary to PWK males, which in the presence of a receptive female always engaged in sexual behavior, a large percentage BL6 males did not become sexual aroused (15 out of 23) and never tried to mount the female (a session was aborted if 30 minutes after female entry the male did not initiate a mount attempt, see Methods for details). Blood was also collected in this condition, at the end of the 30 minutes social interaction (Fig. 1b). In this case, PRL levels of BL6 males did not differ from baseline (baseline 0.99 ± 0.67 vs Social 1.25 ± 0.63; *P* = 0.282, Paired *t* test), further suggesting that PRL is only released in the context of a sexual interaction.

Because PRL is known to be released under stress^46^ and to ensure that the changes observed in circulation are not a result from the blood collection procedure itself, all animals were initially habituated to the collection protocol in another cage, alone. To ensure that the habituation protocol worked, in a separate experiment we measured PRL levels in the absence of any behavior. Four blood samples were collected 20 minutes apart from BL6 and PWK males in their home cage (Fig. 1c). In both cases, circulating PRL levels were not altered, ensuring that the observed increases were not caused by the manipulation (Bl6: F_3,7_ = 2.08; *P* = 0.18; PWK F_3,7_ = 2.94; *P* = 0.11, RM One-way ANOVA).

Collectively, these results demonstrate that PRL is released during sexual behavior in male mice, but not during a social interaction or during the blood collection protocol, prompting us to examine the role of PRL release during sexual behavior.

### Acute prolactin release does not induce a refractory period-like state

To investigate if the increase in circulating levels of PRL that occurs during the sexual interaction is sufficient to decrease sexual motivation, a hallmark of the PERP, we employed a pharmacological approach to acutely elevate PRL levels before the animals became sexually aroused and assess if the male mice behave as if they are in a PERP-like state. PRL is produced in specialized cells of the anterior pituitary, the lactotrophs, and its release is primarily controlled by dopamine originating from the hypothalamus. Dopamine binds D2 receptors at the membrane of the lactotrophs, inhibiting PRL release. Suppression of dopamine discharge leads to disinhibition of lactotrophs, which quickly release PRL into circulation^47,48^. To acutely elevate PRL levels, we performed an intraperitoneal injection of the D2 dopamine receptor antagonist domperidone, which does not cross the blood brain barrier ^49,50^, and measured PRL levels 15 minutes after the procedure. As expected, domperidone administration lead to a sharp rise in the levels of circulating PRL, of similar magnitude to what is observed during copulation (Fig. 2a, BL6: domp 12.54 ± 2.032 vs ejac 7.789 ± 3, *P* = 0.0024; PWK: domp 25.87 ± 7.15 vs ejac 18.55 ± 6.46; *P* = 0.037, Unpaired *t* test).

**Fig. 2.**
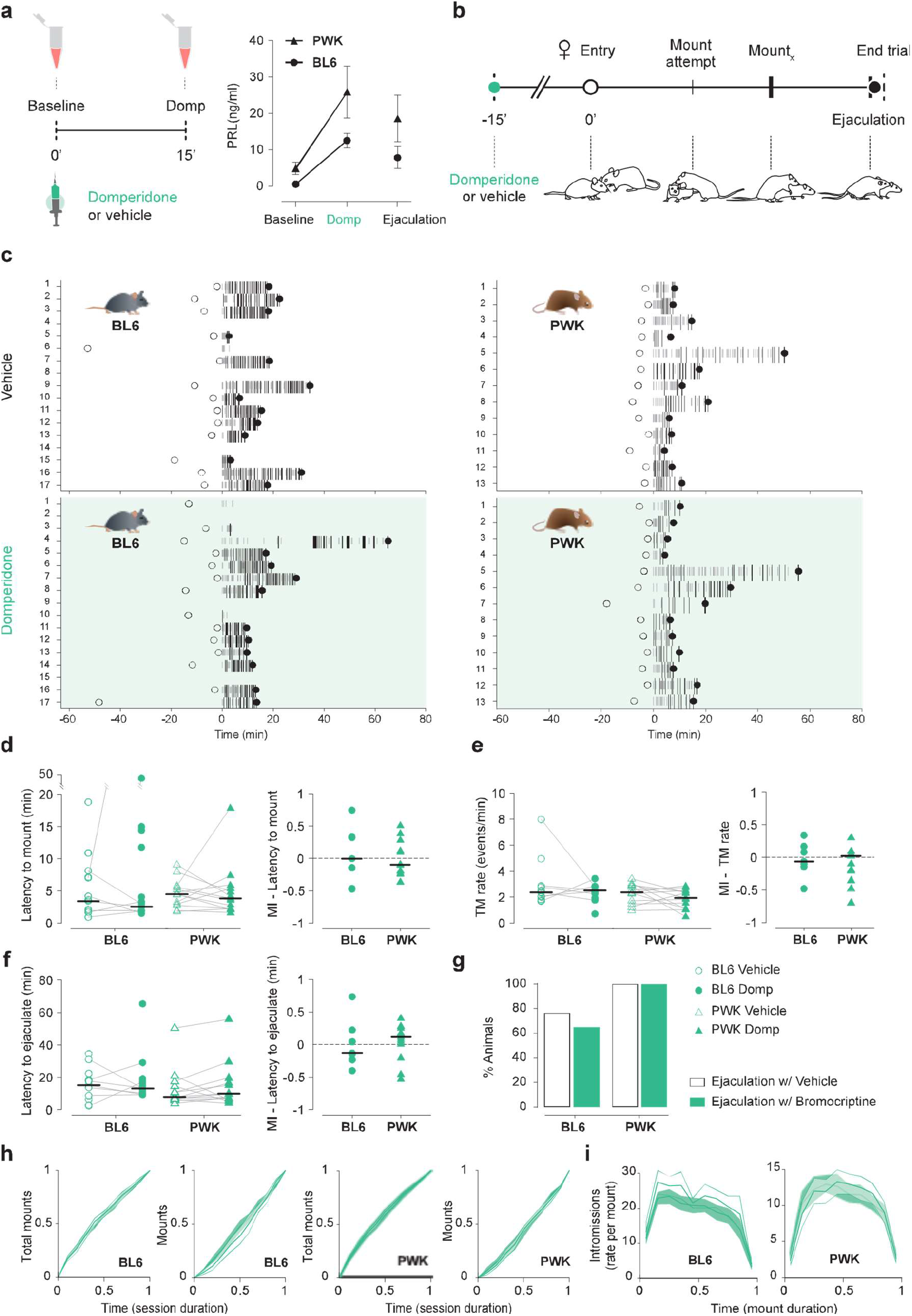
Acute prolactin release does not induce a refractory period-like state. **a** Timeline for blood collection and [PRL]_blood_ after Vehicle (black) or Domperidone (Domp, green) administration. Domp-induced [PRL]_blood_ has a similar magnitude to what is observed during copulation (from Fig. 1 a), two-tailed Unpaired *t* test for BL6 (n = 8) *P* = 0.0024 and PWK (n = 9) *P* = 0.037. **b** Timeline for sexual behavior assay using sexually trained BL6 and PWK males pre-treated with vehicle or Domp (t= −15min). Each animal was tested twice, in a counterbalanced manner: one with vehicle and one with Domp. **c** Raster plot aligned to the first consummately event (first Mount attempt or Mount), representing the sexual behavior executed by the male, with mount attempts represented in small grey bars, mounts in long black bars (width correlated with mount duration) and ejaculation with a black circle. Time of female entry in the apparatus represented with an open circle. BL6 n = 17; PWK n =13. Quantification of **d** Latency to mount (first mount attempt or mount) (BL6 *P* = 1.0, PWK *P* = 0.6355), **e** rate of Total Mounts (TM, mount attempts + mounts) (BL6 *P* = 0.9375, PWK *P* = 0.3054) and **f** Latency to ejaculate (BL6 *P* = 0.8125, PWK *P* = 0.21631). Each line represents data of an individual. Only animals that ejaculated in both sessions were considered in the statistics (n_BL6_ = 7, n_PWK_ = 13). Individuals that did not ejaculate in one of the trials are represented as unconnected dots (not used in statistics). MI [modulation index (domp-veh)/(dom+veh)] between the two conditions for both strains. Data presented as median ± M.A.D. (median absolute deviation with standard scale factor) following Wilcoxon rank sum test. **g** Percentage of animals that reached ejaculation in the vehicle and Domp condition. **h** Cumulative distributions of Total mounts and Mounts along the behavioral assay. Histogram aligned to the first consummatory event with 0,1min bins. Time normalized for the duration of the session from female entry to ejaculation. **j** Rate of intromissions executed during the mount. Histogram for all mounts of each session, aligned to beginning of the mount with 0,1min bins. Time normalized for the duration of the mount.

Therefore, next we investigated how domperidone-treated male mice behave with a receptive female. If PRL is sufficient to induce a PERPlike state, treated males should exhibit decreased sexual motivation, which could be manifested in distinct manners, such as on the latency to initiate consummatory behavior or the vigor of copulation. Each male from the two strains was tested twice, once with vehicle and another time with domperidone, in a counter-balanced manner and all the annotated behaviors are depicted over time on Fig. 2b and c (see Methods for details). Despite differences in the dynamics of sexual behavior across strains, administration of domperidone does not seem to affect sexual motivation, as we could not detect any significant difference in the latency to start mounting the female, frequency of attempts to mount the female, time taken to ejaculate or proportion of animals that reached ejaculation (Fig. 2d-g). Domperidone administration also does not seem to affect the dynamics of the sexual interaction across the session or within each mount (Fig. 2i and j respectively) or other measures of sexual behavioral performance (please see Supplementary Fig. 2).

In summary, domperidone administration, which causes an acute elevation of circulating PRL levels similar to what is observed at the end of copulation, does not have an inhibitory effect on any behavioral parameter related to sexual motivation on the two strains of mice tested, this is, it does not induce a PERP-like state.

### Blocking prolactin release during copulation does not decrease the duration of the refractory period

The release of PRL which is observed during sexual behavior has been proposed to be central in the establishment of the PERP^9^. To test this hypothesis, we acutely inhibited PRL release during sexual behavior by taking advantage of bromocriptine, a D2 receptor agonist. Bromocriptine’s activation of D2 receptors on the lactotrophs’ membrane blocks PRL release, a well-established procedure to inhibit the discharge of this hormone from the pituitary^31,51^. If PRL is indeed necessary for the establishment of the PERP, we expected that after ejaculation, drug-treated males to regain sexual motivation faster than controls.

To test bromocriptine’s efficiency in blocking PRL release during sexual behavior, we first injected males with bromocriptine and measured PRL levels at three times points: i) before the drug or vehicle injection, ii) before the female was inserted in the cage and then iii) after ejaculation (Fig. 3a). As shown in Fig. 3a, bromocriptine administration efficiently blocked PRL release in both subspecies of mice, since PRL levels after ejaculation are not different from baseline (BL6: B1 vs Ejac, *P* =0.3, B2 vs Ejac *P* = 0.99; PWK: B1 vs Ejac, *P* = 0.97, B2 vs Ejac *P* = 0.99; Tukey’s multiple comparisons test after RM Two way Anova).

**Fig. 3.**
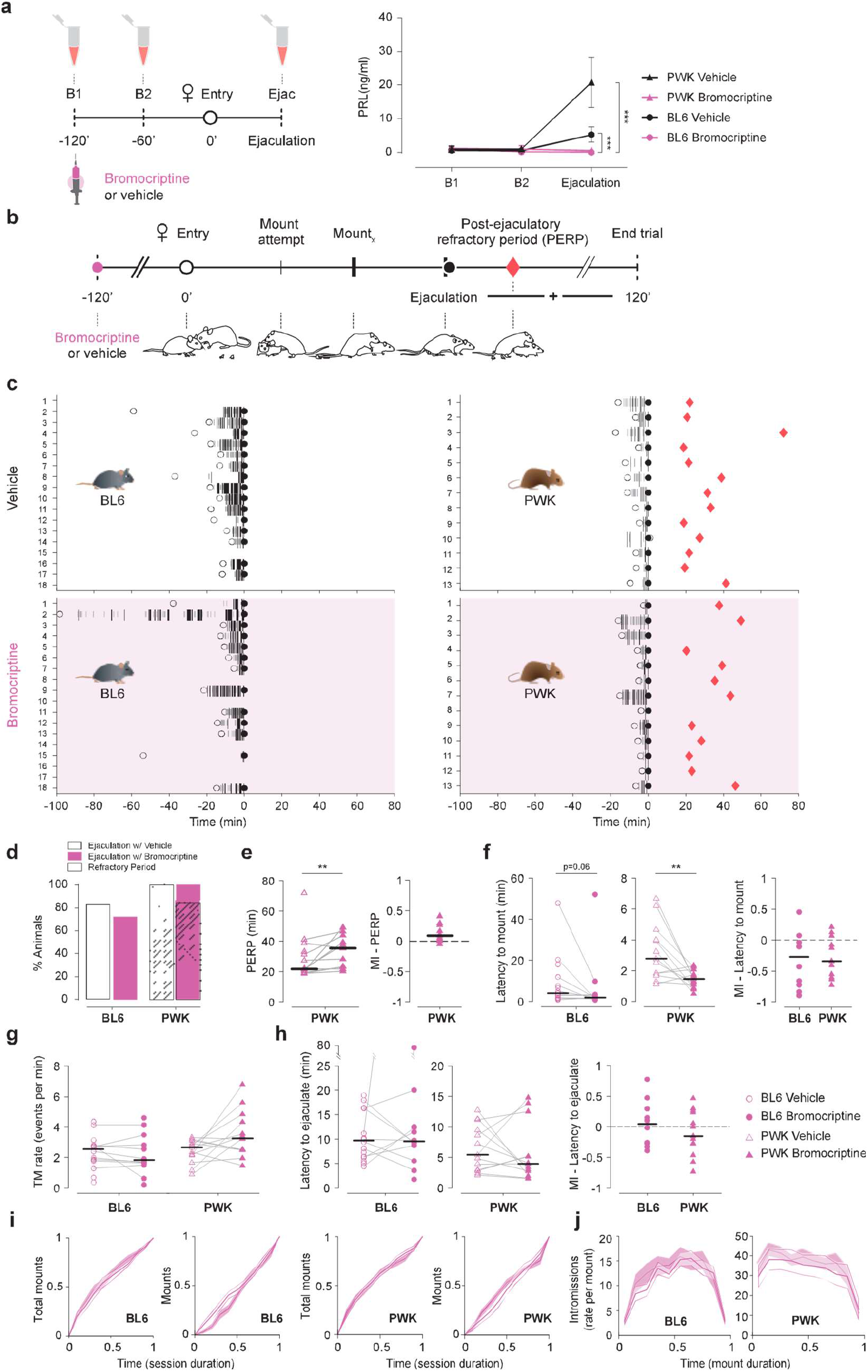
Blocking prolactin release during copulation does not decrease the duration of the refractory period. **a** Timeline for blood collection and [PRL]_blood_ after Vehicle (black) or Bromocriptine (pink) administration. RM two-way Anova with treatment (veh or bromo) as between subject’s factor and time (B1, B2 and Ejac) as the within subject’s factor, followed by Tukey’s multiple comparison test: BL6 (n = 5 each) Treatment F_1,8_=21.08, *P* = 0.0018 Ejac_veh_ vs. Ejac_bromo_ *P* < 0.0001; Time F_2,16_=18.41, *P* < 0.0001, B1_bromo_ vs. Ejac_bromo_ *P* = 0.3; and PWK (n_veh_ = 6, n_bromo_ = 4) Treatment F_1,8_=28.43, *P* = 0.0007 Ejac_veh_ vs. Ejac_bromo_ *P* < 0.0001; Time F_2,16_=23.97, *P* < 0.0001, B1_bromo_ vs. Ejac_bromo_ *P* = 0.97. **b** Timeline for sexual behavior assay using sexually trained BL6 and PWK males pre-treated with vehicle or Bromocriptine (t=-120 min). Each animal was tested twice in a counterbalanced manner. **c** Raster plot aligned to ejaculation, representing the sexual behavior performed by the male, with mount attempts represented in small grey bars, mounts in long black bars (width correlated with mount duration), ejaculation with a black circle and PERP (latency to the first consummatory event (mount attempt or mount) after ejaculation represented with a red diamond. Time of female entry in the apparatus represented with an open circle. BL6 n = 18; PWK, n =13. **d** Percentage of animals that reached ejaculation (solid color) and re-initiated the consummatory behavior after ejaculating (PERP, dashed color) in vehicle (white) and Bromo (pink) conditions. Quantification of **e** PERP duration, **f** latency to mount (first MA or Mount) (BL6 *P* = 0.0644, PWK *P* = 0.0132), **g** rate of Total Mounts (TM, MA+mounts) (BL6 *P* = 0.32, PWK *P* = 0.0681) and **h** latency to ejaculate (BL6 *P* = 0.92, PWK *P* = 0.68). Each line represents data of an individual; Only animals that ejaculated in both sessions were considered in the statistics (n_BL6_ = 10, n_PWK_ = 13). Individuals that did not ejaculate in one of the trials are represented as unconnected dots (not used in statistics). Data presented as median ± M.A.D. (median absolute deviation with standard scale factor) following Wilcoxon rank sum test. MI [modulation index (bromo-veh)/(bromo+veh)] between the two conditions for both strains. \*\*\**P* < 0.0001. **i** Cumulative distributions of Total mounts and Mounts along the behavioral assay. Histogram aligned to the first consummatory event with 0,1min bins. Time normalized for the duration of the session from female entry to ejaculation. **J:** Rate of intromissions executed during the mount. Histogram for all mounts of each session, aligned to beginning of the mount with 0,1min bins. Time normalized for the duration of the mount.

To test the effect of the pharmacological manipulation on PERP duration, the male and female were allowed to remain in the cage undisturbed for a period of up to 2 hours after ejaculation. Each male from the two strains was tested twice, once with vehicle and a second time with bromocriptine, in a counter-balanced manner. Each session ended once the male performed the first attempt of copulation after ejaculation or after two hours if no attempt was made (Fig. 3b, see Methods for details). All the annotated behaviors are depicted over time on Fig. 3b and c (see Methods for details).

As shown in Fig. 3d, inhibiting PRL release during sexual behavior did not change the proportion of male mice of the two strains that reached ejaculation or regained sexual interest in the two hours after ejaculation. Also, and contrary to what was expected, we observed a significant increase in the PERP of PWK males (Fig. 3e, Veh: 21.7 ± 4.18 vs Bromo: 35.4 ± 16.3, *P* = 0.007 by Wilcoxon signed rank test). Administration of bromocriptine seems to affect the initial sexual motivation, as we could detect a decrease in the latency to start mounting the female (Fig. 3f, trend for B6 males and significant for PWK, Veh: 4.06 ± 4.35 vs Bromo: 1.93 ± 1.13, *P* = 0.06; and Veh: 2.78 ± 1.7 vs Bromo: 1.48 ± 0.55, *P* = 0.01, respectively, by Wilcoxon signed rank test). This observation was not due to an increase in activity/locomotion of the bromocriptine treated males as the average male speed before and after the female entry was not affected by the manipulation, nor the distance between the pair. However, besides the locomotor activity being the same, the average male speed projected towards the female increased significantly for PWK treated with bromocriptine as they moved in a goal directed way, directionally towards the female (Supplementary Fig. 3).

However, once consummatory behavior was initiated, control and bromocriptine-treated males exhibited similar levels of sexual motivation, as we could not detect any difference in the frequency of attempts to mount the female or time taken to reach ejaculation (Fig 3g and h). Other aspects of the sexual interaction were also not altered (Supplementary Fig. 3). Furthermore, bromocriptine administration does not seem to affect the dynamics of the sexual interaction across the session or within each mount (Fig. 3i and j respectively).

In summary, blocking PRL release during copulation does not affect the proportion of animals that regain sexual motivation within two hours after ejaculation and contrary to what was expected, bromocriptine leads to an increase in the duration of the PERP of PWK males. Except for a decrease in the latency to start mounting, maintaining circulating PRL low, at levels similar to what is observed prior to the sexual interaction, does not affect any of the parameters of sexual performance analyzed.

## Discussion

The post ejaculatory refractory period or PERP is highly conserved across species and is characterized by a general decrease in sexual motivation after ejaculation^2^. The pituitary hormone PRL is released during copulation and has been put forward as the main player in the establishment of the PERP^9^. However, the involvement of PRL in the establishment and duration of the PERP is controversial and has not been formally tested^2^. Here we show that despite being released during copulation as previously shown in other taxa, PRL is neither sufficient nor necessary for the establishment of the PERP.

In this study we investigated the role of PRL in the PERP of two different strains of mice that belong to the two main subspecies of house mouse, *Mus musculus musculus* (PWK) and *Mus musculus domesticus* (BL6), for two main reasons. As already presented, the two strains have very different PERP duration, widening the dynamic range of this behavioral parameter and increasing the probability of detecting an effect of the manipulations. Second, although fundamental for many present-day discoveries, the usage of the common inbred strains of mice comes at a cost, due to the limitations in their genetic background that sometimes leads to results that are specific to the strain of mouse used^52–54^. Wild derived strains of mice are valuable tools that can complement the genetic deficiencies of classical laboratories strains of mice^44,55,56^. Also, despite the fact that larger numbers of animals are used (because experiments are repeated on each mouse strain), this approach is already routinely used in other fields, such as in immunological studies^57^ providing greater confidence to the results obtained from the effect of pharmacological manipulations on behavior, for example.

We first showed that PRL is released during copulation in male mice. Interestingly, even though being quite an invasive technique, after being habituated to the procedure, sexual behavior does not seem to be affected before or during the consummatory phase. This opens up the possibility to perform such type of experiments using an “within-animal” design, a very important point particularly when there is a large inter-individual variability, while decreasing the number of animals used.

Despite being released during sexual behavior in mice, PRL dynamics are quite different from what has been observed in humans. In men, PRL seems to only be released around the time of ejaculation^15,16^, and only when ejaculation is achieved^16^. Indeed, the fact that PRL surge was only observed when ejaculation was achieved was one of the main results that lead to the idea that PRL may play a role in the acute regulation of sexual arousal after orgasm in humans^58^. In contrast, in mice we observed an increase in circulating levels of PRL in sexually aroused PWK males and in BL6 males during the consummatory phase. The discrepancy between our results and the results published by others might be a result of the sampling procedure. Despite the fact that in human studies blood was continuously collected, PRL detection was performed at fixed time intervals and not upon the occurrence of particular events, such as ejaculation. Therefore, when averaging PRL levels across individuals, each participant might be in a slightly different state of arousal. Also, because PRL concentration is determined over fixed intervals of time, it is difficult to pinpoint the PRL surge to the time of ejaculation (even though the human studies show that sexual arousal per se is not accompanied by an increase in PRL levels)^9^. To our knowledge, a single study assessed PRL levels during sexual behavior in male mice, stating that PRL is released after ejaculation^59^. In this case, blood was also continuously sampled at fixed intervals of time. In contrast, in our study the blood was collected upon the execution of particular events, such as the first mount attempt, a pre-defined number of mounts and ejaculation. Thus, even though the intervals between PRL measurements are different for each mouse, we ensure that PRL levels are measured for all individuals in a similar internal state. Independently of the differences in the dynamics of circulating PRL levels, the raise we observe seems to be specific to a sexual encounter, since PRL levels in BL6 males that never attempt copulation remain unaltered from baseline.

In order to test if PRL by itself is sufficient to decrease sexual motivation, we injected domperidone to induce an artificial PRL-surge. In this case, the male mouse should behave like a male mouse that just ejaculated: for example, exhibit longer latency to initiate the sexual interaction, which in the case of BL6 mice should take days. Even though domperidone administration causes circulating levels of PRL that are similar to the ones observed at the end of a full sexual interaction, this manipulation did not cause any alteration in terms of sexual motivation or performance, as all behavioral parameters remained unaltered for both strains of mice. The fact that, by itself, PRL did not have an impact on sexual motivation might be due to the fact that other neuromodulators and hormones whose levels increase during a normal sexual interaction (serotonin and oxytocin for example)^60^, were not altered by our manipulation. Further experiments could test this idea by examining if combinations of different neuromodulators and hormones administered together can induce a PERP-like state.

Last, we asked if the elevation in PRL levels during sexual behavior is necessary for the establishment and duration of the PERP. For that we took a complementary pharmacological approach, where we injected bromocriptine, a D2 receptor agonist that temporarily inhibits the release of PRL. PRL levels after ejaculation in bromocriptine-treated males are similar to pre-copulatory levels. If PRL is indeed necessary to establish the PERP, we would expect a decrease in its duration that should easily be observed in the PWK males (since they regain sexual interest on average 30 minutes after ejaculation) or even in the BL6 (which take days). The proportion of animals re-engaging in sexual behavior during the 2 hours limit could also be increased for the two strains. We observed a decrease in the latency to start mounting the female and, contrary to our expectation, a significant increase in the PERP duration of PWK males. We believe these effects may be mediated by the direct effect of bromocriptine in the central nervous system, rather than an effect of PRL itself. First, baseline PRL levels are already very low in male mice and therefore the manipulation most likely did not affect them. Second, systemic administration of dopamine agonists has shown that anticipatory measures of sexual behavior are more sensitive to disruption than are consummatory measures of copulation^61,62^. This agrees with our results, where we observed a significant decrease in the latency to initiate mounting with bromocriptine, while no other parameter of sexual performance was affected. Interestingly, bromocriptine-treated PWK males seem more ballistic in their approach to the female, suggesting a more goal-directed behavior towards the female. Bromocriptine (and domperidone) might also have an effect outside the central nervous system as D2 receptors are expressed in the human and rat seminal vesicles^63^. It is not known if direct manipulation of these receptors in the seminal vesicles has an impact on the PERP.

What could be the role of copulatory PRL? PRL release may be the “side-effect” of the neuromodulatory changes that occur during sexual behavior, this is, merely the result of reduction in DA levels (DA inhibits PRL release) and/or the increase in oxytocin and serotonin (known stimulating factors of PRL release) instead of having the principal role in the establishment of PERP^64–66^. The fact that PRL levels are already elevated during the sexual interaction in BL6 and PWK males, further suggests that PRL cannot promote by itself reduced sexual motivation, at least in male mice. Other studies point towards a role of PRL in the establishment of parental behavior^67,68^. New behavioral paradigms will be fundamental aid in unravelling this mystery.

## Methods

### Animals

BL6 (*Mus musclus domesticus*, C57BL/6J) and Wild (*Mus musculus musculus*, PWD/PhJ and PWK/PhJ) mice were ordered from The Jackson Laboratories and maintained in our animal facility. Animals were weaned at 21 days and housed in same-sex groups in stand-alone cages (1284L, Techniplast, 365 x 207 x 140 mm) with access to food and water ad libitum. Mice were maintained on a 12:12 light/dark cycle and experiments were performed during the dark phase of the cycle, under red dim light. All experiments were approved by the Animal Care and Users Committee of the Champalimaud Neuroscience Program and the Portuguese National Authority for Animal Health (Direcção Geral de Veterinária).

Females were kept house grouped and males were isolated before the sexual training. Both males and females were sexually experienced. Males interacted with different females in each sexual encounter. Animals were habituated to be handled and to the assay routine to reduce stress. All experiments were conducted in parallel for both BL6 and PWK. Trials were conducted in the male home cage (1145T, Techniplast, 369 x 156 x 132 mm) striped from nesting, food and water; covered with a transparent acrylic lid. The trial started with the entry of the female in the setup (t=0min).

### Ovariectomy and hormonal priming

All females underwent bilateral ovariectomy under isoflurane anesthesia (1-2% at 1L/min). After exposing the muscle with one small dorsal incision (1 cm) a small incision was made in the muscle wall, at the ovary level, on each side. The ovarian arteries were cauterized and both ovaries were removed. The skin was sutured, and the suture topped with iodine and wound powder. The animals received an ip injection of carpofen before being housed individually with food supplemented with analgesic (MediGel, 1mg carprofen /2 oz cup) for 2 days recovery and then re-grouped in their home cages.

Female mice were primed subcutaneously 48 hours before the assay with 0,1ml estrogen (1mg/ml, Sigma E815 in sesame oil) and 4 hours before the assay with 0,1ml progesterone (5mg/ml, Sigma 088K0671 in sesame oil).

### Blood collection

Tail-tip whole blood sampling was done as previously described^45^. Briefly, blood was collected from the male tail, immediately diluted in PBS-T (PBS, 0.05% Tween20) and frozen at −20°C straightaway, where it was stored until use.

To profile [PRL]_blood_ during sexual behavior (Fig. 1a), baseline blood was collected 30 minutes before (t=-30min) the female entry (t=0min). From this point on, blood collection was locked to the onset of specific behaviors: once the male did the first mount attempt (MA), after executing of a fixed number of mounts (Mx) and after ejaculation. We choose Mx=5 for BL6 and Mx=3 for PWK to ensure that the males would have significant sexual interaction without reaching ejaculation. Contrarily to PWK males that, in the presence of a receptive female, the majority engages in sexual behavior, BL6 do not. Thus, after 30 minutes interacting with the female without displaying sexual interest, we collected a blood sample and terminated the trail (Fig. 1b, social). Because blood collection is an invasive procedure and PRL is also released under stress we evaluated if the manipulation itself could induce PRL release. For that we collected blood every 20 minutes for 1 hour from males resting in their home cage (Fig. 1c).

Domperidone is a d2 antagonist that was previously used to study the inhibitory tone of dopamine on PRL release from the pituitary, inducing a PRL peak 15 minutes after ip injection ^45^. To test the magnitude of the PRL release of the two mouse strains under domperidone (Fig 2a), we conducted a pilot study where we collected a blood sample before (baseline) and 15 minutes after domp injection (20mg/kg, abcam Biochemicals). We opted to manipulate [PRL]_blood_ trough domperidone instead of injecting PRL directly to induce a PRL release similarly to a natural occurring instead of adding a recombinant form.

Bromocriptine is a D2 dopamine receptor agonist known to inhibit endogenous prolactin release^31^. To test its efficacy on blocking PRL release during sexual behavior (Fig. 3a) we conducted a second pilot study where the males were injected (100 μg bromo or vehicle) 2 hours before the trial started. Blood samples were collected just before injection, 1 hour after injection and after ejaculation.

### Prolactin quantification

[PRL]_blood_ quantification was done as previously described^45^. Briefly, a 96-well plate (Sigma-Aldrich cls 9018–100EA) was coated with 50 μl capture antibody antirat PRL (anti-rPRL-IC) (National Institute of Diabetes and Digestive and Kidney Diseases (NIDDK), AFP65191 (Guinea Pig), NIDDK-National Hormone and Pituitary Program (NHPP, TORRANCE, CA) at a final dilution of 1:1000 in PBS of the antibody stock solution, reconstituted in PBS as described in the datasheet (Na2HPO4 7.6 mM; NaH2PO4 2.7 mM and NaCl 0.15M; pH 7.4). The plate was protected with Parafilm^®^ and incubated at 4°C overnight in a humidified chamber. The coating antibody was decanted and 200 μl of blocking buffer (5% skimmed milk powder in PBS-T) was added to each well to block nonspecific binding. The plate was left for 2 hours at room temperature on a microplate shaker. In parallel, a standard curve was prepared using a 2-fold serial dilution of Recombinant mouse Prolactin (mPRL; AFP-405C, NIDDK-NHPP) in PBS-T with BSA 0.2 mg/mL (bovine serum albumin; Millipore 82–045–1). After the blocking step, the plate was washed (3 times for 3 minutes at room temperature with PBS-T), 50μl of quality control (QC), standards or samples were loaded in duplicate into the wells and incubated for 2 hours at room temperature on the microplate shaker. The plate was washed, and the complex was incubated for another 90 minutes with 50 μl detection antibody (rabbit alpha mouse PRL; a gift from Patrice Mollard Lab) at a final dilution of 1:50 000 in blocking buffer solution. Following a final wash, this complex was incubated for 90 minutes with 50 μl horseradish peroxidase-conjugated antibody (anti rabbit, IgG, Fisher Scientific; NA934) diluted in 50% PBS, 50% blocking buffer. One tablet of O-phenylenediamine (Life technologies SAS 00–2003) was diluted into 12 ml Citrate-phosphate buffer pH 5, containing 0.03% hydrogen peroxide. 100 μl of this substrate solution was added to each well (protected from light), and the reaction was stopped after 30 minutes with 50 μl of 3M HCl. The optical density from each well was determined at 490nm using a microplate reader (SPECTROstar^Nano^, BMG LABTECH). An absorbance at 650nm was used for background correction.

A linear regression was used to fit the optical densities of the standard curve vs their concentration using samples ranging from 0.1172ng/ml to 1.875ng/ml. Appropriate sample dilutions were carried out in order to maintain detection in the linear part of the standard curve. PRL concentrations were extrapolated from the OD of each sample. To control for reproducibility of the assay, trunk blood of males injected with domperidone was immediately diluted in PBS-T and pulled to be used as quality control (QC). Loading of the wells was done vertically left to right and QC was always loaded on the top row. The formula OD (Co,t) = OD (Ob) + α (QC).t was used to correct the ODs for loading dwell time (OD: optical density, Co: corrected, t: well number, Ob: observed, α: QC linear regression’ α). Coefficient of variability was kept to a maximum of 10%.

### Behavioral assays

Each male underwent two trials: one with vehicle and one with drug (domperidone or bromocriptine). Administrations were counter balanced between animals and spaced seven days. In the first assay, for pharmacological induction of acute PRL release (Fig. 2b), the male was injected ip with domperidone or vehicle 15 minutes before the trial started (t= −15min). Animals were allowed to interact until the male reached ejaculation or 1 hour in the case the male did not display sexual behavior. Conversely, for pharmacological blockage of PRL release (Fig. 3b), a second group of males were pre-treated with bromocriptine or vehicle with a subcutaneous injection 2 hours before the beginning of the trial (t=-120min). Animals were allowed to interact until a maximum of 2 hours after the male reached ejaculation or 1h in the case the male did not display sexual behavior.

### Behavior analysis

The behavior was recorded from the top and side with pointgrey cameras (FL3-U3-13S2C-CS) connected to a computer running a custom Bonsai software^69^. Behavior was manually annotated using the open source program Python Video Annotator (https://pythonvideoannotator.readthedocs.io) and analyzed using Matlab. The number of mount attempts (MA, mount without intromission), mounts (mounts with intromission), latency to mount (first MA or mount), latency to ejaculation and PERP (latency between ejaculation and the next mount) was calculated. Total number of mounts (TM) was calculated as the sum of MA and mounts and TM rate was calculated as (TM)/(latency to ejaculate). The percentage of animals that reached ejaculation and regain sexual interest under 2 hours (Refractory period) were also calculated. The modulation index (MI) was calculated as (X_drug_-X_vehicle_)/(X_drug_+X_vehicle_). The centroid position and individual identity of each pair was followed off-line using the open source program idtracker.ai^70^ and used to calculate male velocity and inter individual distance with Matlab (Supplementary Fig. 3)

### Statistical analysis

The statistical details of each experiment, including the statistical tests used and exact value of n are detailed in each figure legend. Data related to prolactin quantification was analyzed using GraphPad Prism 7 software and presented as mean ± S.D. For comparison within strain (Fig. 1a and c) an RM One-way Anova followed by a Tukey’s multiple comparison test was used. Comparison of paired samples comparing two groups, statistical analysis was performed by using a paired-sample two-tailed *t* test (Fig. 1b, 2a baseline-Domp and 3a). Analysis between unpaired samples comparing two groups was performed using an unpaired-sample two-tailed *t* test (Fig. 2a Domp-Ejaculation). Data related to animal behavior was analyzed with MATLAB R2019b and presented as median ± M.A.D. (median absolute deviation with standard scale factor). Animals were randomized between treatments and comparison between the two conditions were done with Wilcoxon rank sum test (Fig. 2d-f, 3e-h and supplementary Fig. 4 to 6). Only animals that ejaculates in both session were included in the statistical comparisons (Domperidone: n_BL6_ = 7, n_PWK_ = 13; Bromocriptine: n_BL6_ = 10, n_PWK_ = 13). Significance was accepted at *P* < 0.05 for all tests.

## Supporting information

Supp Valente et al

## Data availability

All data generated to support the findings of this study are available from the corresponding author upon reasonable request.

## Acknowledgements

We thank Patrice Mollard for welcoming S.V. in his laboratory to learn the uElisa technique and for provision of antibodies. We also express our gratitude to Francisco Romero for all the support with IdTracker and to Gil Costa for the figure design. This work was supported by Champalimaud Foundation, ERC consolidator Grant (772827), Fundação para a Ciência e Tecnologia SFRH / BD / 51011 / 2010 (S.V.) and PhRMA Foundation Postdoctoral Fellowship in Informatics (T.M).

## Autor contribution

S.Q.L. and S.V. designed the study. S.V. did the experiments, annotation of the behavior and IdTracker. T.M. wrote the Matlab code. S.V. analyzed the data with input from S.Q.L. and T.G. S.Q.L. and S.V. wrote the paper with contributions from others.

## Competing interests

The authors declare no competing interests.

## Footnotes

* https://en.wikipedia.org/wiki/Refractory_period; https://www.humanitas.net/treatments/prolactin

## Notes

### Competing Interest Statement

The authors have declared no competing interest.

## References

1. Masters, W. H. & Johnson, V. E. Human sexual response. (Little Brown, Boston, 1966).

2. Seizert, C. A. The Neurobiology of the Male Sexual Refractory Period. Neurosci Biobehav Rev 92, 350–377 (2018).

3. Levin, R. J. Revisiting Post-Ejaculation Refractory Time—What We Know and What We Do Not Know in Males and in Females. J Sex Medicine 6, 2376–2389 (2009).

4. McGill, T. E. Sexual Behavior of the Mouse after Long-Term and Short-Term Postejaculatory Recovery Periods. J Genetic Psychology 103, 53–57 (1963).

5. Rodriguez-Manzo, G. Blockade of the establishment of the sexual inhibition resulting from sexual exhaustion by the Coolidge effect. Behav Brain Res 100, 245–254 (1999).

6. Wilson, J. R., Kuehn, R. E. & Beach, F. A. Modification in the sexual behavior of male rats produced by changing the stimulus female. J Comp Physiol Psych 56, 636 (1963).

7. Rojas-Durán, F. et al. Correlation of prolactin levels and PRL-receptor expression with Stat and Mapk cell signaling in the prostate of long-term sexually active rats. Physiol Behav 138, 188–192 (2015).

8. Hernandez, M. E. et al. Prostate response to prolactin in sexually active male rats. Reprod Biol Endocrin 4, 28 (2006).

9. Kruger, T. H. C., Haake, P., Hartmann, U., Schedlowski, M. & Exton, M. S. Orgasm-induced prolactin secretion: feedback control of sexual drive? Neurosci Biobehav Rev 26, 31–44 (2002).

10. Drago, F. Prolactin and sexual behavior: A review. Neurosci Biobehav Rev 8, 433–439 (1984).

11. Freeman, M. E., Kanyicska, B., Lerant, A. & Nagy, G. Prolactin: Structure, Function, and Regulation of Secretion. Physiol Rev 80, 1523–1631 (2000).

12. Grattan, D. R. & Kokay, I. C. Prolactin: A Pleiotropic Neuroendocrine Hormone. J Neuroendocrinol 20, 752–763 (2008).

13. Brody, S. & Krüger, T. H. C. The post-orgasmic prolactin increase following intercourse is greater than following masturbation and suggests greater satiety. Biol Psychol 71, 312–315 (2006).

14. Egli, M., Leeners, B. & Kruger, T. H. C. Prolactin secretion patterns: basic mechanisms and clinical implications for reproduction. Reproduction Camb Engl 140, 643–654 (2010).

15. Exton, M. S. et al. Coitus-induced orgasm stimulates prolactin secretion in healthy subjects. Psychoneuroendocrino 26, 287–294 (2001).

16. Exton, N. G. et al. Neuroendocrine response to film-induced sexual arousal in men and women. Psychoneuroendocrino 25, 187–199 (2000).

17. Krüger, T. et al. NEUROENDOCRINE AND CARDIOVASCULAR RESPONSE TO SEXUAL AROUSAL AND ORGASM IN MEN. Psychoneuroendocrino 23, 401–411 (1998).

18. Krüger, T. H. C. et al. Serial neurochemical measurement of cerebrospinal fluid during the human sexual response cycle. Eur J Neurosci 24, 3445–3452 (2006).

19. Exton, M. S. et al. Cardiovascular and Endocrine Alterations After Masturbation-Induced Orgasm in Women. Psychosom Med 61, 280–289 (1999).

20. Kruger, T. et al. Specificity of the neuroendocrine response to orgasm during sexual arousal in men. J Endocrinol 177, 57–64 (2003).

21. Oaknin, S., Castillo, A. R. D., Guerra, M., Battaner, E. & Mas, M. Changes in forebrain Na,K-ATPase activity and serum hormone levels during sexual behavior in male rats. Physiol Behav 45, 407–410 (1989).

22. Haake, P. et al. Absence of orgasm-induced prolactin secretion in a healthy multi-orgasmic male subject. Int J Impot Res 14, 3900823 (2002).

23. Svare, B. et al. Hyperprolactinemia Suppresses Copulatory Behavior in Male Rats and Mice. Biol Reprod 21, 529–535 (1979).

24. Buvat, J. Hyperprolactinemia and sexual function in men: a short review. Int J Impot Res 15, 3901043 (2003).

25. Sato, F. et al. Suppressive Effects of Chronic Hyperprolactinemia on Penile Erection and Yawning Following Administration of Apomorphine to Pituitary-Transplanted Rats. J Androl 18, 21–25 (1997).

26. Melmed, S. et al. Diagnosis and Treatment of Hyperprolactinemia: An Endocrine Society Clinical Practice Guideline. J Clin Endocrinol Metabolism 96, 273–288 (2011).

27. Grattan, D. R. & Bridges, R. S. Hormones, Brain and Behavior (Second Edition). Endocr Syst Interact Brain Behav 2471–2504 (2009) doi:10.1016/b978-008088783-8.00079-6.

28. Riddle, O., Bates, R. W. & Dykshorn, S. W. THE PREPARATION, IDENTIFICATION AND ASSAY OF PROLACTIN—A HORMONE OF THE ANTERIOR PITUITARY. Am J Physiology-legacy Content 105, 191–216 (1933).

29. Ganong, W. F. Circumventricular Organs: Definition And Role In The Regulation Of Endocrine And Autonomic Function. Clin Exp Pharmacol P 27, 422–427 (2000).

30. Grattan, D. R. et al. Feedback Regulation of PRL Secretion Is Mediated by the Transcription Factor, Signal Transducer, and Activator of Transcription 5b. Endocrinology 142, 3935–3940 (2001).

31. Brown, R. S. E., Kokay, I. C., Herbison, A. E. & Grattan, D. R. Distribution of prolactin-responsive neurons in the mouse forebrain. J Comp Neurology 518, 92–102 (2010).

32. Salais-López, H., Agustín-Pavón, C., Lanuza, E. & Martínez-García, F. The maternal hormone in the male brain: Sexually dimorphic distribution of prolactin signaling in the mouse brain. Plos One 13, e0208960 (2018).

33. Esteves, F. F., Matias, D., Mendes, A. R., Lacoste, B. & Lima, S. Q. Sexually dimorphic neuronal inputs to the neuroendocrine dopaminergic system governing prolactin release. J Neuroendocrinol 31, (2019).

34. Kirk, S. E., Grattan, D. R. & Bunn, S. J. The median eminence detects and responds to circulating prolactin in the male mouse. J Neuroendocrinol 31, e12733 (2019).

35. Georgescu, T., Ladyman, S. R., Brown, R. S. E. & Grattan, D. R. Acute effects of prolactin on hypothalamic prolactin receptor expressing neurones in the mouse. Biorxiv 2020.05.10.087650 (2020) doi:10.1101/2020.05.10.087650.

36. Bole-Feysot, C., Goffin, V., Edery, M., Binart, N. & Kelly, P. A. Prolactin (PRL) and Its Receptor: Actions, Signal Transduction Pathways and Phenotypes Observed in PRL Receptor Knockout Mice. Endocr Rev 19, 225–268 (1998).

37. Corona, G., Jannini, E. A., Vignozzi, L., Rastrelli, G. & Maggi, M. The hormonal control of ejaculation. Nat Rev Urology 9, 508 (2012).

38. Levin, R. Is prolactin the biological “off switch” for human sexual arousal? SexRelatsh Ther 18, 237–243 (2003).

39. Turley, K. R. & Rowland, D. L. Evolving ideas about the male refractory period. Bju Int 112, 442–452 (2013).

40. Kamel, F., Right, W. W., Mock, E. J. & Frankel, A. I. The Influence of Mating and Related Stimuli on Plasma Levels of Luteinizing Hormone, Follicle Stimulating Hormone, Prolactin, and Testosterone in the Male Rat 1. Endocrinology 101, 421–429 (1977).

41. Kamel, F. & Frankel, A. J. Hormone Release during Mating in the Male Rat: Time Course, Relation to Sexual Behavior, and Interaction with Handling Procedures*. Endocrinology 103, 2172–2179 (1978).

42. Kruger, T. et al. Effects of acute prolactin manipulation on sexual drive and function in males. J Endocrinol 179, 357–365 (2003).

43. Lenschow, C. & Lima, S. Q. In the mood for sex: neural circuits for reproduction. Curr Opin Neurobiol 60, 155–168 (2020).

44. Gregorová, S. & Forejt, J. PWD/Ph and PWK/Ph inbred mouse strains of Mus m. musculus subspecies--a valuable resource of phenotypic variations and genomic polymorphisms. Folia Biol-prague 46, 31–41 (2000).

45. Guillou, A. et al. Assessment of Lactotroph Axis Functionality in Mice: Longitudinal Monitoring of PRL Secretion by Ultrasensitive-ELISA. Endocrinology 156, 1924–1930 (2015).

46. Gala, R. R. The physiology and mechanisms of the stress-induced changes in prolactin secretion in the rat. Life Sci 46, 1407–1420 (1990).

47. Lyons, D. J. & Broberger, C. TIDAL WAVES: Network mechanisms in the neuroendocrine control of prolactin release. Front Neuroendocrin 35, 420–438 (2014).

48. Grattan, D. R. 60 YEARS OF NEUROENDOCRINOLOGY: The hypothalamo-prolactin axis. J Endocrinol 226, T101–T122 (2015).

49. Cocchi, D. et al. Prolactin-Releasing Effect of a Novel Anti-Dopaminergic Drug, Domperidone, in the Rat. Neuroendocrinology 30, 65–69 (1980).

50. Costall, B., Fortune, D. H. & Naylor, R. J. Neuropharmacological studies on the neuroleptic potential of domperidone (R33812). J Pharm Pharmacol 31, 344–347 (1979).

51. Molik, E. & Błasiak, M. The Role of Melatonin and Bromocriptine in the Regulation of Prolactin Secretion in Animals – A Review. Ann Anim Sci 15, 849–860 (2015).

52. puglisi-Allegra, S. & Cabib, S. PSYCHOPHARMACOLOGY OF DOPAMINE: THE CONTRIBUTION OF COMPARATIVE STUDIES IN INBRED STRAINS OF MICE. Prog Neurobiol 51, 637–661 (1997).

53. Cabib, S., Puglisi-Allegra, S. & Ventura, R. The contribution of comparative studies in inbred strains of mice to the understanding of the hyperactive phenotype. Behav Brain Res 130, 103–109 (2002).

54. Welker, E. & Loos, H. V. der. Quantitative correlation between barrel-field size and the sensory innervation of the whiskerpad: a comparative study in six strains of mice bred for different patterns of mystacial vibrissae. J Neurosci 6, 3355–3373 (1986).

55. Fernandes, C. et al. Behavioral Characterization of Wild Derived Male Mice (Mus musculus musculus) of the PWD/Ph Inbred Strain: High Exploration Compared to C57BL/6J. Behav Genet 34, 621–630 (2004).

56. Onos, K. D. et al. Enhancing face validity of mouse models of Alzheimer’s disease with natural genetic variation. Plos Genet 15, e1008155 (2019).

57. Deschepper, C. F., Olson, J. L., Otis, M. & Gallo-Payet, N. Characterization of blood pressure and morphological traits in cardiovascular-related organs in 13 different inbred mouse strains. JApplPhysiol 97, 369–376 (2004).

58. Krüger, T. H. C., Hartmann, U. & Schedlowski, M. Prolactinergic and dopaminergic mechanisms underlying sexual arousal and orgasm in humans. World J Urol 23, 130–138 (2005).

59. Bronson, F. H. & Desjardins, C. Endocrine Responses to Sexual Arousal in Male Mice*. Endocrinology 111, 1286–1291 (1982).

60. Pfaus, J. G. Pathways of Sexual Desire. J Sex Medicine 6, 1506–1533 (2009).

61. Pfaus, J. G. & Phillips, A. G. Role of Dopamine in Anticipatory and Consummatory Aspects of Sexual Behavior in the Male Rat. Behav Neurosci 105, 727–743 (1991).

62. Hull, E. M., Muschamp, J. W. & Sato, S. Dopamine and serotonin: influences on male sexual behavior. Physiol Behav 83, 291–307 (2004).

63. Hyun, J.-S. et al. Localization of Peripheral Dopamine D1 and D2 Receptors in Rat and Human Seminal Vesicles. J Androl 23, 114–120 (2002).

64. McIntosh, T. K. & Barfield, R. J. Brain monoaminergic control of male reproductive behavior. I. Serotonin and the post-ejaculatory refractory period. Behav Brain Res 12, 255–265 (1984).

65. McIntosh, T. K. & Barfield, R. J. Brain monoaminergic control of male reproductive behavior. II. Dopamine and the post-ejaculatory refractory period. Behav Brain Res 12, 267–273 (1984).

66. Carmichael, M. S. et al. Plasma Oxytocin Increases in the Human Sexual Response*. J Clin Endocrinol Metabolism 64, 27–31 (1987).

67. Brown, R. E. Hormones and Paternal Behavior in Vertebrates. Am Zool 25, 895–910 (1985).

68. Schradin, C. & Anzenberger, G. Prolactin, the Hormone of Paternity. Physiology 14, 223–231 (1999).

69. Lopes, G. et al. Bonsai: an event-based framework for processing and controlling data streams. Front Neuroinform 9, 7 (2015).

70. Romero-Ferrero, F., Bergomi, M. G., Hinz, R. C., Heras, F. J. H. & Polavieja, G. G. de. idtracker.ai: tracking all individuals in small or large collectives of unmarked animals. Nat Methods 16, 179–182 (2019).

