## Supplementary material for "No evidence for prolactin’s involvement in the post-ejaculatory refractory period": Supp Valente et al

**Supplementary Fig. 1**

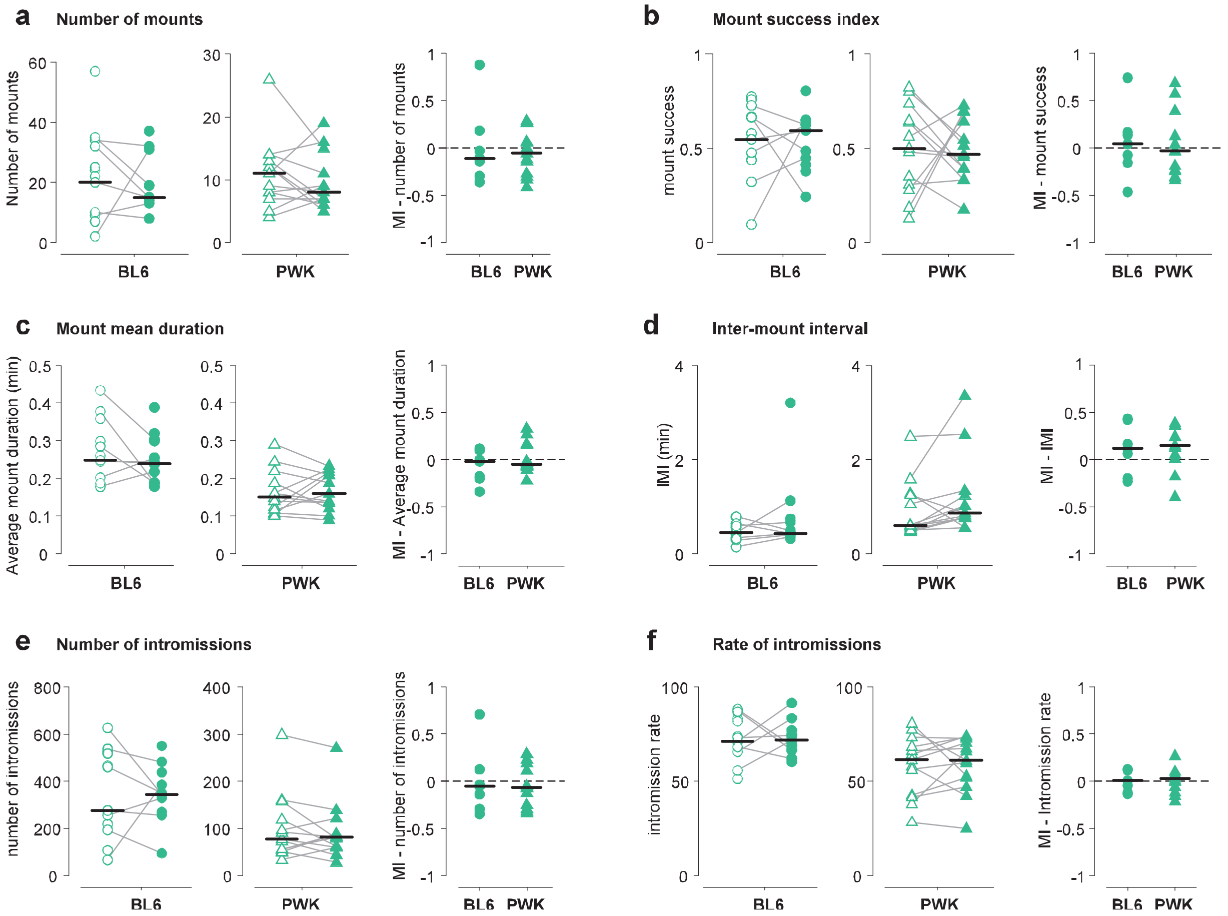


**Supplementary Fig. 1** Additional analysis of the sexual behavior assay using adult sexually trained BL6 and PWK males pre-treated with vehicle or Domperidone (total number of subjects used: BL6 n = 17; PWK, n =13; with ejaculation in both sessions BL6 n = 7; PWK, n =13 ). Quantification of **a** number of mounts, **b** mount success index [mounts/(mount attempt + mounts)], **c** mount mean duration (total mount duration/#mounts), **d** Inter-mount interval (IMI), **e** number of intromissions and **f** rate of intromissions (#intromissions/total mount duration). Each animal underwent two behavioral assays: one with vehicle and one with domperidone. Administration was counterbalanced between animals. Each line represents data of an individual; Unconnected dots represent data of an individual that did not ejaculate in the matching trial (data not used for statistics). MI [modulation index (domp-veh)/(dom+veh)]. Data presented as median ± M.A.D. (median absolute deviation with standard scale factor) following Wilcoxon rank sum test.

**Supplementary Fig. 2**

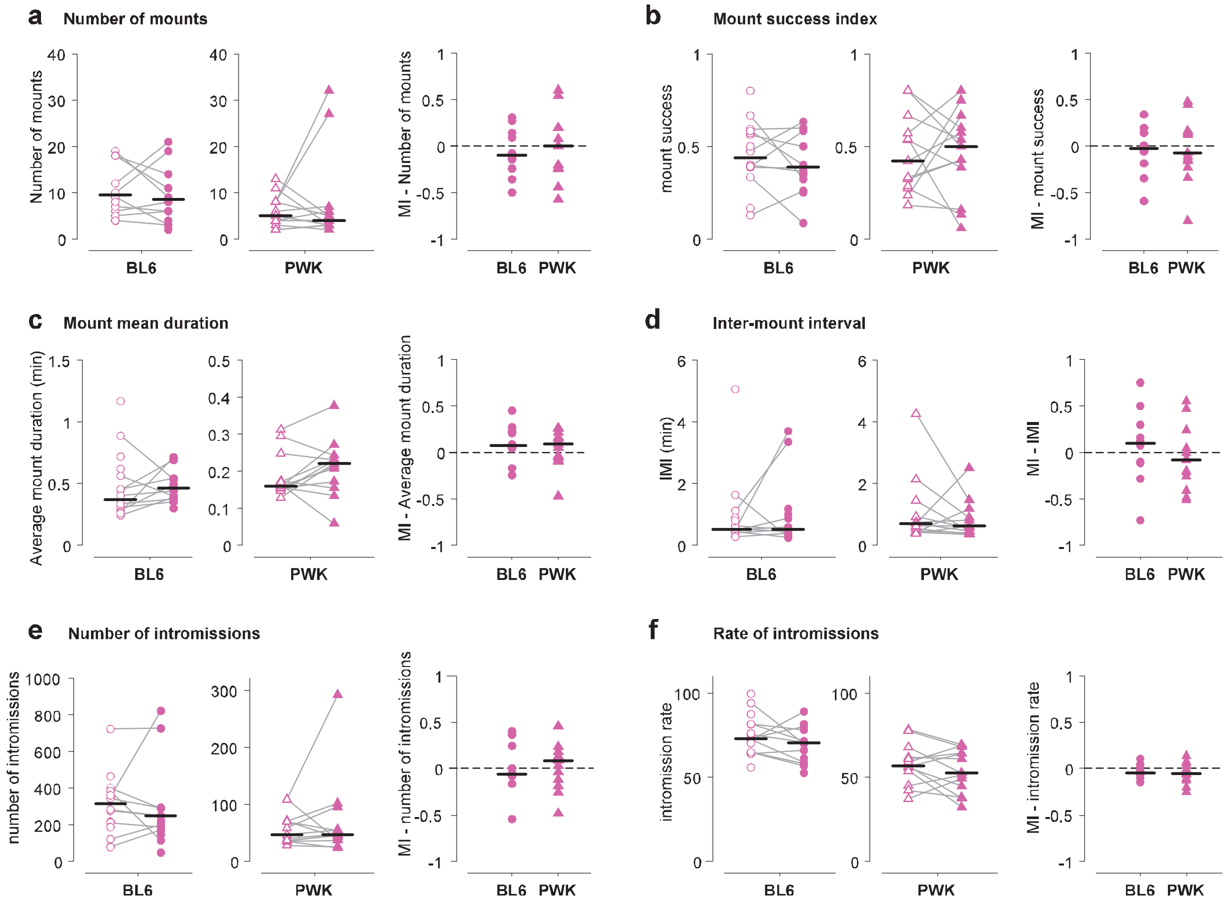


**Supplementary Fig. 2** Additional analysis of the sexual behavior assay using adult sexually trained BL6 and PWK males pre-treated with vehicle or Bromocriptine (total number of subjects used: BL6 n = 18; PWK, n =13; with ejaculation in both sessions BL6 n = 10; PWK, n =13). Quantification of **a** number of mounts, **b** mount success index [mounts/(mount attempt + mounts)], **c** mount mean duration (total mount duration/#mounts), **d** Inter-mount interval (IMI), **e** number of intromissions and **f** rate of intromissions (#intromissions/total mount duration). Each animal underwent two behavioral assays: one with vehicle and one with domperidone. Administrations were contra balanced between animals. Each line represents data of an individual; Unconnected dots represent data of an individual that did not ejaculate in the matching trial (data not used for statistics). MI [modulation index (bromo-veh)/(bromo+veh)]. Data presented as median ± M.A.D. (median absolute deviation with standard scale factor) following Wilcoxon rank sum test. Significance was accepted at $P < 0.05$.

**Supplementary Fig. 3**

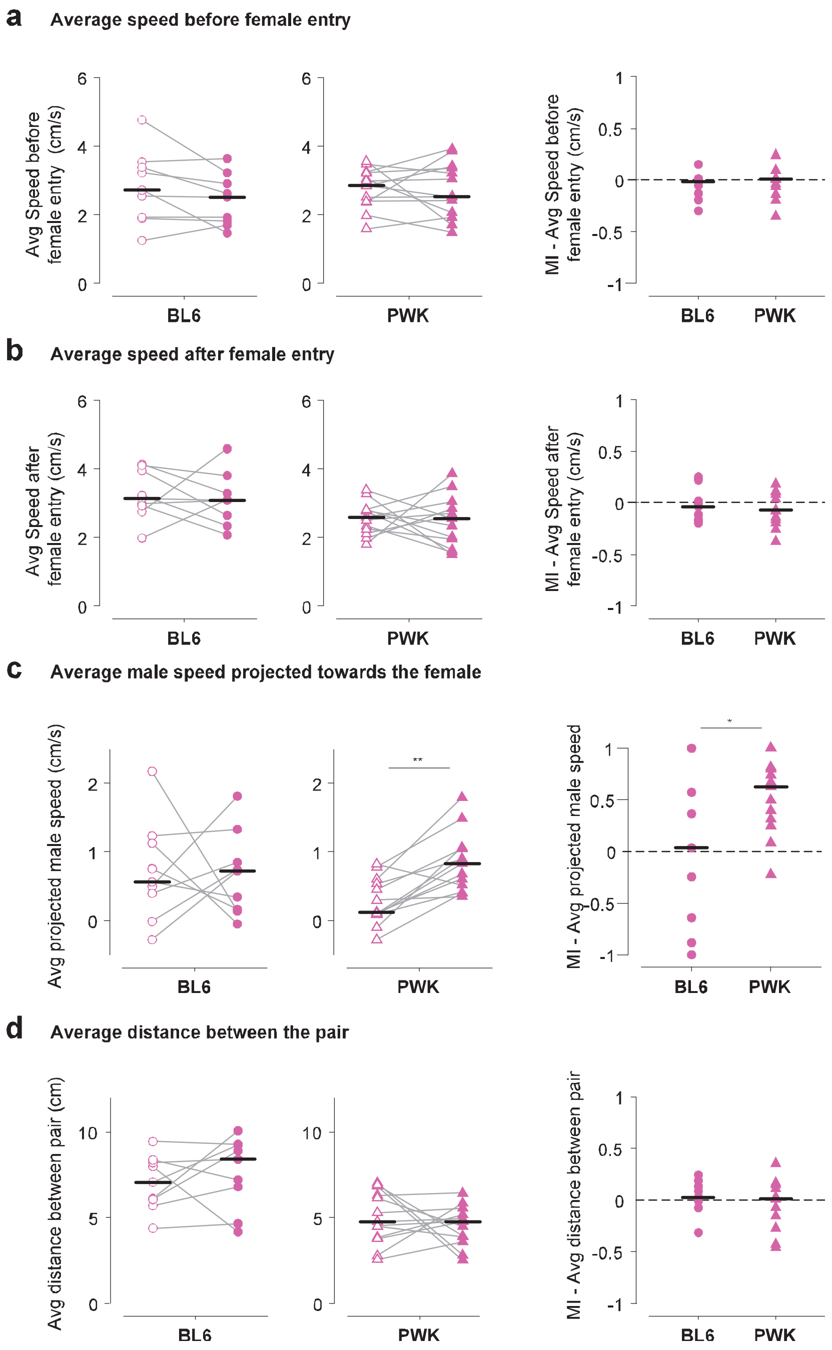


**Supplementary Fig. 3** Bromocriptine treated animals no not show increased locomotion activity. Average speed of the male **a** before and **b** after the female entry until the first consummatory event. **c** Average male speed projected towards the female **d** average distance between the pair. Metrics were taken from couples that reached ejaculation in both session from bromocriptine assay BL6 n= 9, PWK n =13 (Fig. 3). Each line represents data of an individual; MI [modulation index (bromo-veh)/(bromo+veh)]. Data presented as median ± M.A.D. (median absolute deviation with standard scale factor) following Wilcoxon rank sum test. Significance was accepted at $P < 0.05$. ** $P < 0.001$, * $P < 0.01$.
